# Harnessing the power of regional baselines for broad-scale genetic stock identification: A multistage, integrated, and cost-effective approach

**DOI:** 10.1101/2023.03.20.533554

**Authors:** Bobby Hsu, Christopher Habicht

## Abstract

Genetic stock identification (GSI) estimates the contribution of each population to a mixture and these analyses are usually conducted regionally using genetic baselines specific to the stocks expected in that region. Often these regional baselines cannot be combined to produce broader geographical baselines. In cases where the mixture contains stocks spanning across a wide area a broad-scale baseline is created, but these baselines often are unable to resolve among regional stocks. Here, we introduce a new GSI method to harness the resolution capabilities of baselines developed for regional applications in the analyses of mixtures containing fish from a broad geographic range. This multistage process allows for disparate baselines to be used in a single integrated process that estimates the propagated errors from each stage. The baselines used by this model do not require any overlap in markers or in populations representing the broad-scale or regional baselines. The integrated multistage framework allows GSI of a wide geographic area without first developing a large scale, high resolution genetic baseline, or dividing a mixture sample into smaller regions beforehand. This approach is more cost-effective than updating range-wide baselines with all critical regionally important markers.

## Introduction

Genetic stock identification (GSI) is used to estimate the stock composition of fishery mixtures by comparing mixture genotypes to a baseline of allele frequency estimates from populations that represent all stocks (reporting groups) that may be present in the mixture. This method requires that the baseline includes allele frequency estimates for all the genetic markers for all the populations. Genetic baselines are often developed regionally, specifically designed to distinguish among local stocks in regional fishery mixtures (e.g Flannery et al. 2010; Ackerman et al. 2011; Ensing et al. 2013; Gilbey et al. 2018; Beacham et al. 2020; Euclide et al. 2021; Beacham et al. 2021). Often these regional baselines cannot easily be combined across regions to produce broader geographical baselines because the type of genetic markers (e.g., microsatellites, single nucleotide polymorphisms) do not match or do not overlap at an adequate set of loci.

Traditionally, researchers from multiple laboratories have screened common sets of markers across populations representing larger geographic areas to conduct broad-scale GSI analyses, such as GSI of high-seas mixtures (e.g., Beacham et al. 2005; Beacham et al. 2009; McKinney et al. 2020, Wirgin et al. 2020). These baselines require a high level of coordination among laboratories to standardize genetic markers designed to allocate fish to broad-scale reporting groups, and they have been used successfully (Stephenson et al. 2009; Seeb et al. 2007). However, increasing resolution at the regional level can be expensive and logistically challenging because additional markers may be necessary to sufficiently resolve fine-scale reporting groups. Incorporating new or additional genetic markers into a single broad-scale baseline can require multiple laboratories to genotype their respective baseline populations or extensively share their baseline tissues or DNA with the laboratory doing the baseline update. Obtaining tissues or DNA from the required coastwide populations can be difficult, especially if tissue samples are unavailable because they have been exhausted in previous analyses or are not shared by other researchers. Genotyping the large numbers of fish in broad scale baselines is also costly.

As an alternative, some researchers have adopted a “hard cut-off” two-step GSI approach: fish are first individually assigned to broad-scale reporting groups using established coastwide genetic baselines, then, fish assigned to certain reporting groups with probabilities surpassing a set threshold are genotyped for additional genetic markers and assigned to fine-scale reporting groups using a regional baseline (e.g., Miller et al. 2010; Samarasin et al. 2019). However, in this approach, these two GSI analyses are independent to each other such that individual fish might be misassigned in the first step, and the uncertainty incurred cannot be accounted for in the second step.

Here, we developed an integrated multistage GSI model that combines the two-step method into a single process. Our multistage model is essentially two Bayesian GSI models (Pella-Masuda model hereafter; Pella & Masuda 2001) stacked on top of each other; hence the name “multistage.” Estimations of population compositions in a Bayesian GSI model is done through iterations of Markov Chain Monte Carlo algorithm called the Gibbs sampler (Casella & George 1992). We constructed our multistage GSI framework by incorporating the two-step method in each iteration of the Gibbs sampler process. The multistage workflow starts with a mixture sample with each fish genotyped for all the genetic markers in both the broad-scale and regional baselines. During each iteration of the Gibbs sampler, each fish is assigned to a population in the broad-scale baseline in the first stage of the analysis. And at the second stage, each fish is assigned to a population using the regional baseline, but only fish that are assigned to populations within regional baseline reporting groups during the first stage will be counted at the second stage. In other words, assignment of the regional groups is based on assignment of the broad-scale groups. Reporting group proportions are estimated at each stage and combined at the end of each iteration based on this conditional relationship. This algorithm is repeated thousands of times until the estimates converged into a stable distribution (i.e., estimates stabilized). The two stages are integrated as a single process that holistically encapsulates errors incurred at both stages.

We demonstrate our integrated multistage framework using a case study, with mixture data synthesized from existing genetic baselines using a “leave-one-out” analysis. That is, genotypes of fish are randomly selected from existing baselines to synthesize a mixture data set, and the fish selected for the mixture are excluded from the baseline data sets. Because we set the group proportions of the simulated data set, we can quantify the precision and accuracy of the estimates by comparing them to the true values. Also, using the results of leave-one-out analysis, we compared integrated multistage method to the conventional hard cut-off two-step approach.

## Methods

### The model

We constructed our multistage model by placing two Pella-Masuda models hierarchically as integrated stages. The two stages required five pieces of information to estimate reporting group proportions in a mixture sample: a) allele frequencies for each population in a baseline that is capable of distinguishing among broad-scale reporting groups, b) allele frequencies for each population in a baseline (not necessarily the same or overlapping markers used in the broad-scale baseline) that is capable of distinguishing among within-region reporting groups, c) the grouping of populations into reporting groups for both baselines, d) prior information about the reporting group proportions for both baselines, and e) the genotypes of fish sampled. To initiate the Gibbs sampler, each individual in the mixture sample is assigned an initial population identity based on the five pieces of information. Number of individuals in each population is summarized to estimate initial population proportions. Once the initial estimates are obtained, the Gibbs sampler proceeds in the following steps:

1. All individuals in the mixture sample are assigned to new populations and sorted into reporting groups in the broad-scale baseline based on information in a), c), d) and e) and broad-scale population proportions estimated in the previous iteration (first iteration uses the initial estimates).
2. All individuals are assigned to populations and sorted into reporting groups in the regional baseline based on information in b), c), d) and e) and regional population proportions estimated in the previous iteration (first iteration uses the initial estimates). Fish that are assigned to reporting groups outside of the regional baselines during the first stage are ignored.
3. Both broad-scale and regional population proportions are estimated based on the results of individual assignments at each stage.
4. The estimates are updated by repeating steps 1 through 3 until the estimates stabilized.

We summarized the integrated multistage genetic stock identification workflows in Figure 1.

**Figure 1:**
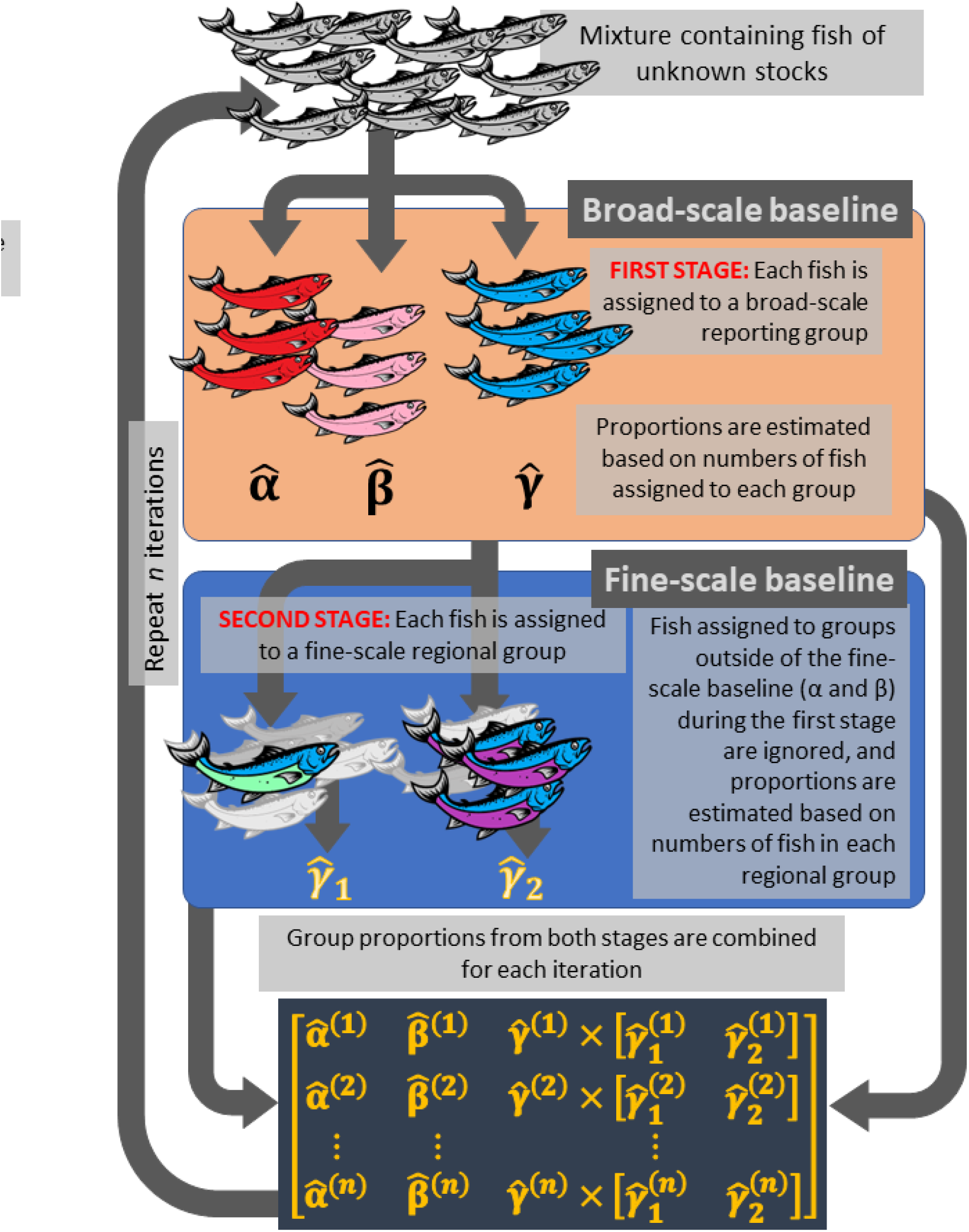
Summary of workflows for genetic stock identification in an integrated multistage framework. Steps are repeated *n* iterations, and *n* should be large enough so that the estimates converge into a stable distribution. Reporting group proportions estimated at each stage were combined and recorded during each iteration. At the first stage, the broad-scale reporting groups were *A, B*, and *C* with respective proportions *α, β*, and *γ*. Using a regional baseline, *C* was further divided into regional reporting groups *C*_1_ and *C*_2_ at the second stage with respective proportions *γ*_1_ and *γ*_2_. We combined the group proportions from the two stages as follows: *α, β, γ* · (*γ*_1_, *γ*_2_).

In the integrated multistage framework, the two Pella-Masuda models are connected because a fish being assigned to a regional group at the second stage is conditional on whether that fish is assigned to a reporting group within region of the regional baseline during the first stage. We modify slightly the notations in Pella & Masuda (2001) to describe the conditional relationship between the broad-scale and the regional baselines:

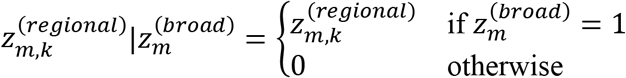

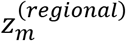 are vectors of indicators (0 or 1) identifying the regional population that individual *m* belongs to. 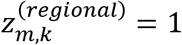 if individual *m* belongs to population *k* and 0 otherwise. 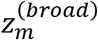 is an indicator as well, and 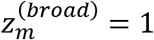 if individual *m* belongs to a reporting group within the regional baseline and 0 otherwise. Combining *z*^(*regional*)^ and 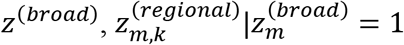 if individual *m* belongs to a reporting group within the regional baseline and is assigned to population *k* in the regional baseline.

Individually, vectors 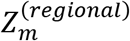 have a multinomial distribution, 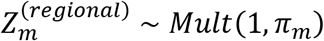, where *π_m_* are vectors of probabilities for individual *m* being assigned to each regional reporting group. The prior distribution for *π_m_* is a Dirichlet, *Dirich*(*K*^-1^), where *K* is the number of reporting groups in the regional baseline. We refer to Pella & Masuda (2001) for detailed descriptions of the posterior distribution of *Z*.

In the context of the integrated multistage framework, whether an individual is assigned to a within-region reporting group is decided during the first stage (i.e. using the broad-scale baseline); therefore, individual identity has a binomial outcome at the first stage and is expressed as 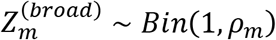. The prior for *ρ_m_* has a Beta distribution with two scale parameters; one equals to the proportions of broad-scale stocks that are within-region of the regional baseline, and the other equals to the proportions of broad-scale stocks that are outside of the regional baseline. The expectation for *ρ_m_* equals to the proportion of broad-scale stocks within the regional baseline.

Probability of an individual being assigned to regional population *k* can be derived using the Bayes rule:

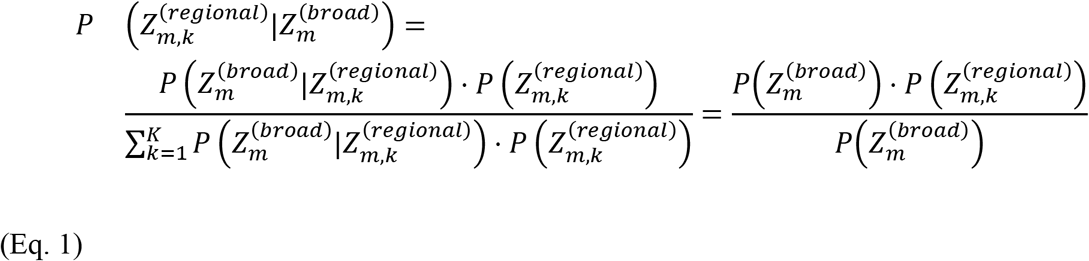

Equation 1 can be simplified into 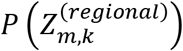 because the outcome of *z*^(*broad*)^ does not depend on the outcome of *z*^(*regional*)^.

We modified the Pella-Masuda model by adapting the “conditional Bayesian GSI” approach (Moran & Anderson 2018) in our Gibbs sampler algorithm. We programmed the Gibbs sampler in *R* (R Core Team, 2022) and is available freely as a *R* package, Ms.GSI, at https://github.com/boppingshoe/Ms.GSI. *R* code and the components of the package can be viewed at the same Github site.

### Case Study

We demonstrate the utility of our integrated multistage model (Ms.GSI) by comparing it to the hard cut-off two-step method (HC 2-step) and the true proportions of a simulated mixture. We used a simulated mixture sample assembled from existing Chinook salmon (*Oncorhynchus tshawytscha*) baselines for the Northern Bering Sea, Alaska. The DNA samples were collected during various projects in the past, and their genotype data were archived in a database maintained by the Alaska Department of Fish & Game Gene Conservation Lab (GCL). We used a marker set of 380 loci that could effectively differentiate Chinook salmon among four reporting groups from the middle/upper Yukon River (Koyukuk, Tanana, Upper U.S., and Canada; Lee et al. 2021; Figure 2C). We retrieved genotypes from the GCL database for this set of genetic markers from fish collected at 151 spawning sites for populations across Yukon River and coastal western Alaska (Coastal W. Alaska; Figure 2A). We synthesized a mixture sample containing an arbitrary number of 205 fish from populations from all five reporting groups from the coastal western region of Alaska and the Yukon River (Figure 2A) in proportions shown in Table 1.

**Figure 2:**
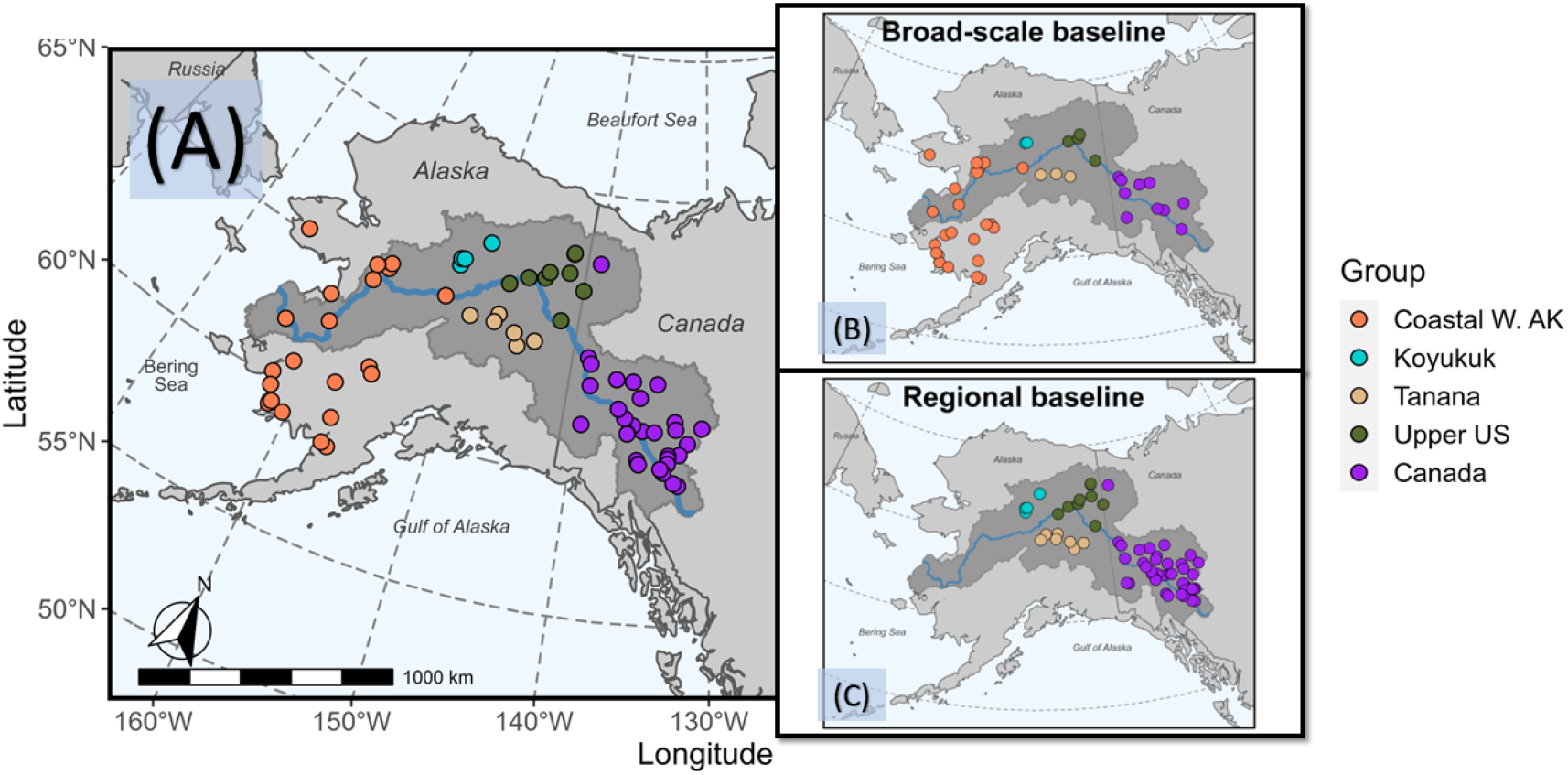
Map showing the collection locations in the North Bering Sea case study for A) the simulated mixture sample, B) the broad-scale baseline (10 markers), and C) the regional baseline (380 markers). Fill color represents reporting groups of Chinook salmon populations. Shaded area represents the Yukon River drainage.

**Table 1:**
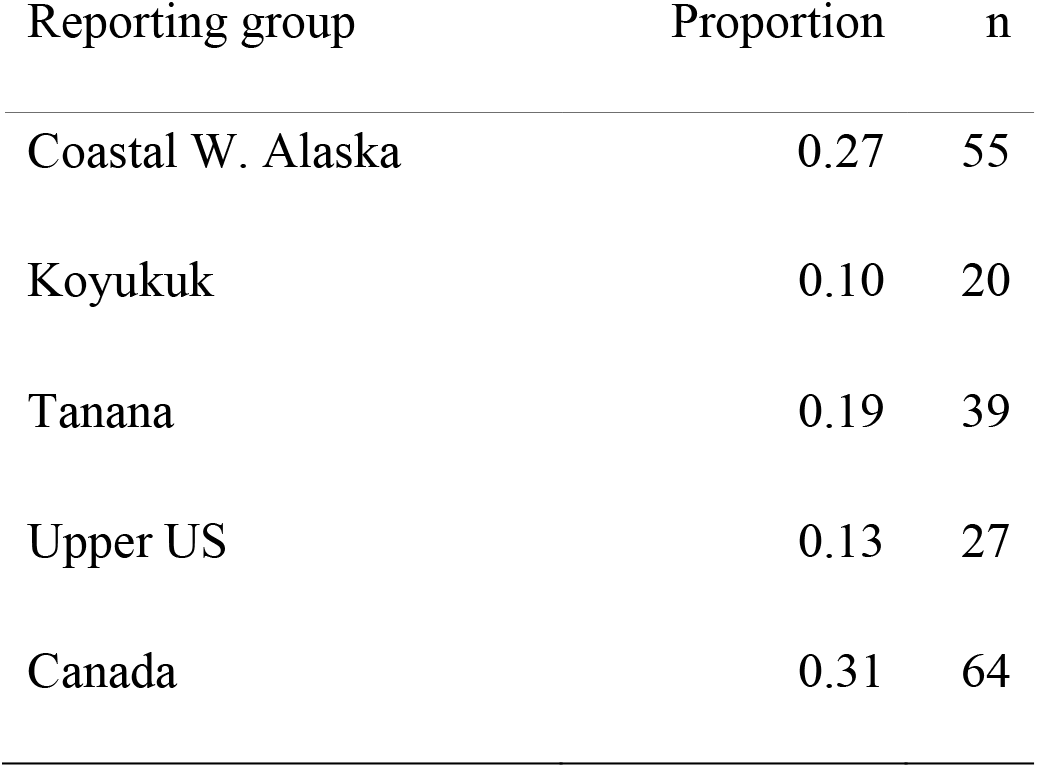
Reporting group proportions and sample sizes for synthetic mixtures in the leave-one-out analysis for Bering Sea Chinook.

We used the baseline for the Northern Bering Sea juvenile Chinook salmon (84 loci and 111 collections; Howard et al. 2019; Figure 2B) as the broad-scale baseline. We included only a subset of the genetic markers screened in order to simulate a depauperate baseline that was capable distinguishing among broad-scale reporting groups, but not the fine-scale reporting groups. We reduced the broad-scale baseline data set to include only 10 of these genetic markers for all populations, which was adequate to distinguish two broad-scale reporting groups (Coastal W. Alaska and middle/upper Yukon River). These 10 loci were also a subset of the 380 marker-set described in the previous paragraph. For the regional baseline, we used the full set of 380 genetic markers, but pared down the populations to only those representing the middle/upper Yukon River reporting groups (106 collections; Figure 2C). And this baseline was able to distinguish among the fine-scale reporting groups. Fish selected for the mixture data set were excluded from both baseline data sets.

### Ms.GSI

We conducted the analysis and ran the Gibbs sampler for the integrated multistage model using our own *R* package, Ms.GSI. To run the Gibbs sampler, we set the prior probabilities for all reporting groups to be equal for each baseline. We ran the sampler in five independent chains, each with 25,000 iterations. We discarded the first 15,000 iterations of each chain as warm-up runs (i.e., burn-ins) and thinned the resulting posterior distribution by keeping every fifth sequence. We checked for convergence of the estimates among chains using Gelman-Rubin diagnostics and effective size (Gelman et al. 2014; Gelman & Rubin 1992) and by visually inspecting the trace plots for amount of mixing between chains (Brooks & Gelman 1998).

### HC 2-step

We also conducted a GSI analysis on the same data set using a conventional hard cut-off two-step approach. We first assigned each fish in the data set to a broad-scale reporting group. Fish that were assigned to a reporting group in the regional baseline with a probability greater than 0.8 were selected for the second stage. We use the regional baseline for the second GSI analysis. We conducted GSI for each baseline separately, using the Gibbs sampler script we developed in *R* and followed the same procedures for the integrated multistage model to run the Gibbs sampler.

### Model comparisons

We compared the results of Ms.GSI and the hard cut-off two-step approach to the true proportions of the synthesized data. We summarized and graphically inspected the assignment probabilities at the regional level.

To quantify the performance of the models, we followed the procedures for genetic baseline evaluation outlined in Barclay et al. (2019) with modifications. We repeated the leave-one-out analyses 50 more times, each time with randomly chosen group proportions for the synthesized mixture sample. To shorten computing time for the evaluation, we ran the 50 leave-one-out analyses each with five chains of 5,000 iterations instead of 25,000. Because posterior means were the only metric needed for the model evaluation, simulations with five chains of 5,000 iterations would satisfy such purpose using the data sets we had. We discarded the first half of each chain as warm-up run and did not thin the posterior output.

We compared the mean estimates and the true values of the group proportions for each of the 50 analyses. To measure how often the estimates deviate from the true values, we calculated the portions of mean estimates that were allocated beyond ± 0.1 of the true values of the mixture proportions. Furthermore, we calculated the root mean squared errors (RMSE) of the mean estimates for each model. Root mean squared error quantifies bias and precision, and a smaller value for RMSE indicates less bias and higher precision. It is calculated as 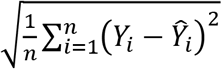, where *n* is number of analyses (50 in this case), *Y_i_* is the true proportion of group *y* for the *i*th iteration, and 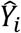 is the mean estimated proportion of group *y* for the *i*th iteration.

The 50 leave-one-out assessments also served as an evaluation to assess the adequacy of the broad-scale baselines we selected for our analyses. Because the power of the baselines was decreased with a reduced marker set, we made sure that the reduced baselines could adequately distinguish the broad-scale reporting groups chosen for our analyses. We followed the evaluation guidelines described in Barclay et al. (2019). Mainly, the estimates should fall within ± 0.1 of the true values more than 90% of the time and have RMSE ≤ 0.05. For our analysis, only broad-scale baseline needs to be assessed.

## Results

For the single leave-one-out analysis, convergence diagnostics showed no major concerns for the Gibbs sampler simulations. Baseline evaluation showed that the reduced broad-scale baseline had an acceptable resolution in identifying broad-scale reporting groups, albeit RMSE was slightly above the guideline. Tables for estimates summary, convergence diagnostics, and model evaluations can be found in Appendix A.

Hard cut-off two-step approach rendered estimates for the four regional groups only, while Ms.GSI provided estimates for all five groups. Ms.GSI showed slightly larger credible intervals for their estimates of the regional groups compared to the hard cut-off two-step approach (Figure 3).

**Figure 3:**
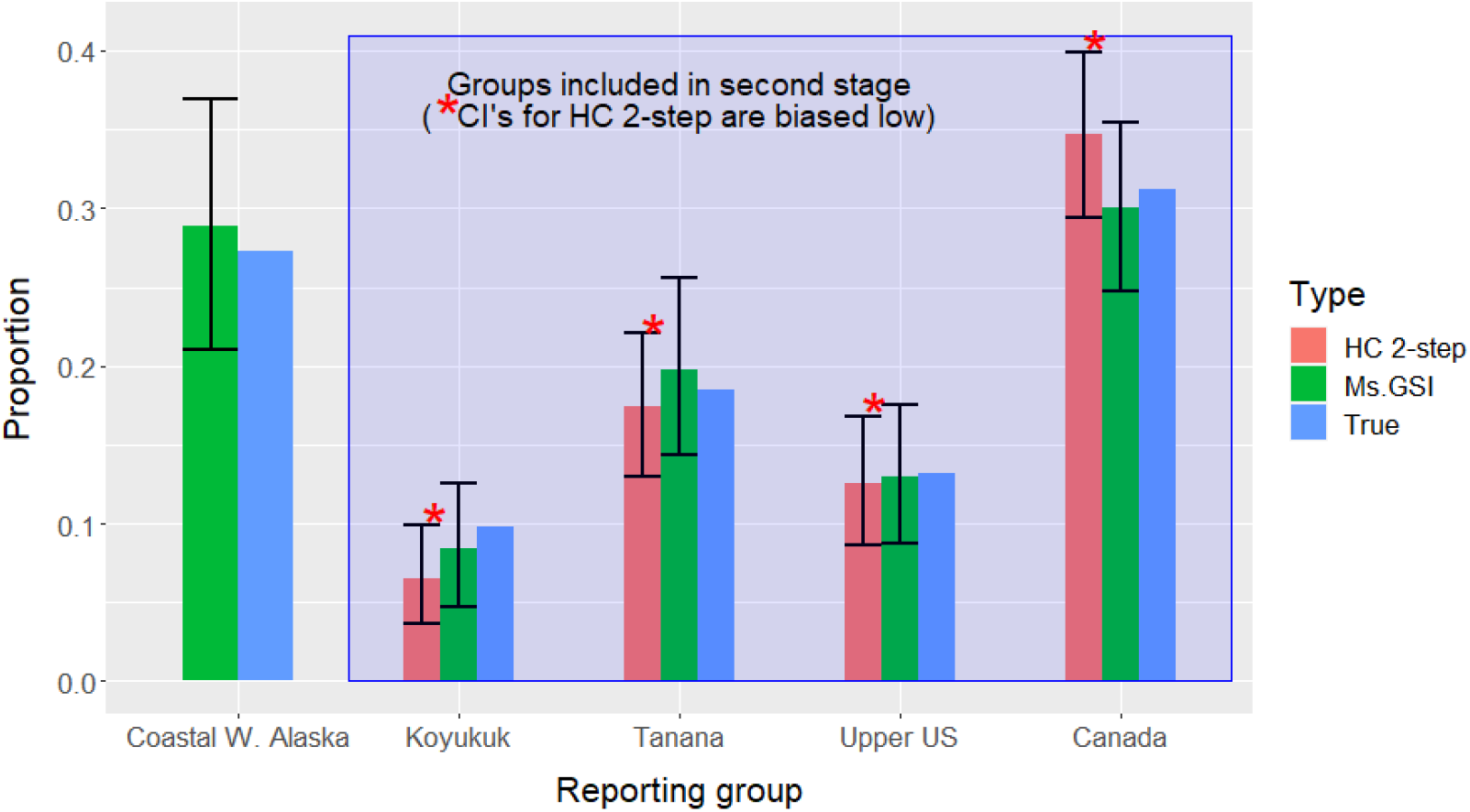
Comparison between estimates of integrated multistage framework (Ms.GSI) and the hard cut-off two-step approach (HC 2-step) and true proportions of the five reporting groups in the leave-one-out analysis for Northern Bering Sea Chinook.

Among the regional reporting groups, Koyukuk had the lowest average probability to be assigned to the second stage and was the most likely to be misassigned compared to other Yukon groups (Tanana, Upper US, and Canada; Figure 4).

**Figure 4:**
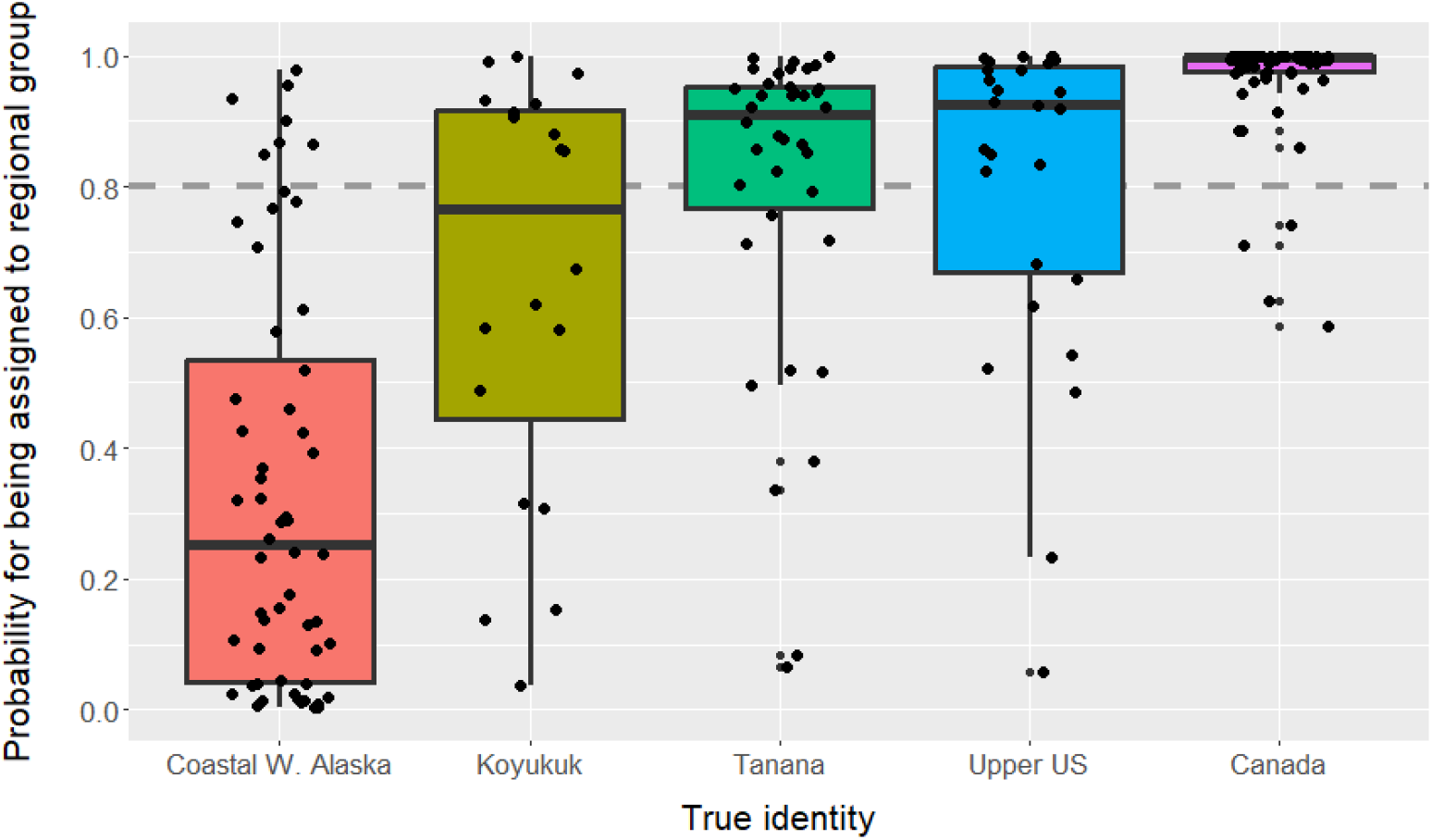
Individual assignment probabilities for assigning fish from each reporting group (x-axis) to the middle/upper Yukon River region during the first stage of the analysis. The true identities of the reporting groups are represented by different colors. Each dot represents an individual fish in the data. The grey dashed line indicates the arbitrarily set 0.8 probability threshold for individual reporting group assignment.

Model evaluation based on 50 leave-one-out analyses indicated that the integrated multistage framework improved estimates by lowering deviation and RMSE compared to a hard cut-off two-step GSI model (Figure 5; Appendix A).

**Figure 5:**
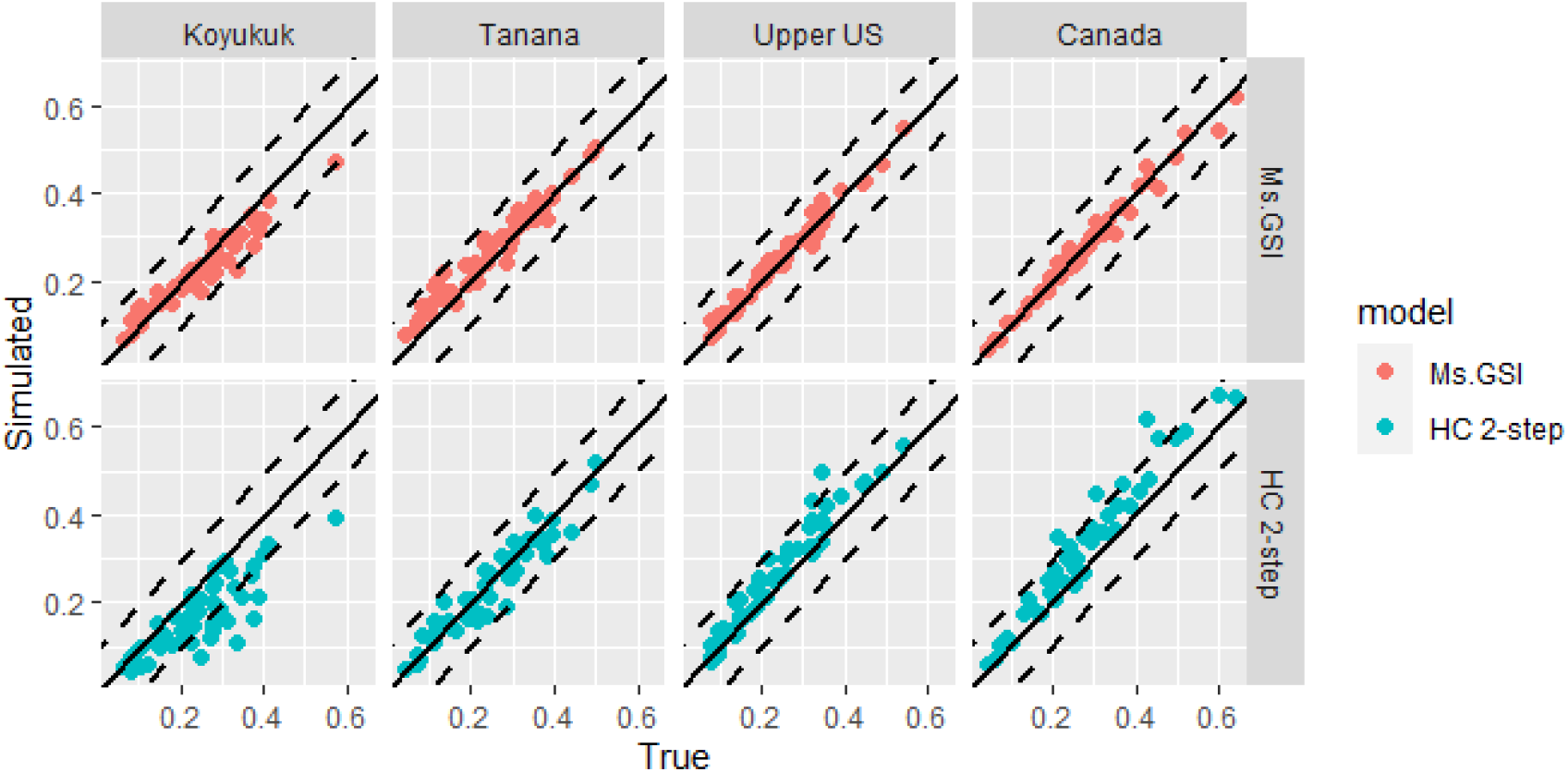
Mean estimated proportions are plotted against the true values for each of the 50 leave-one-out analyses. The solid diagonal lines represent where the estimate equals the true value. The dashed lines represent ± 0.1 of the true value. All methods use Bering Sea Chinook baseline as the broad-scale baseline and Yukon River Chinook baseline as the regional baseline. Coastal Western Alaska group is excluded from this plot because it does not go through the second stage of the analysis for the two-step approaches.

## Discussion

We introduced and tested an integrated multistage GSI method that can harness the fine-scale resolution capabilities of regional baselines in the analyses of mixtures containing fish from a broad geographic range. This integrated multistage framework allows for disparate baselines to be used in a single integrated process that properly propagates errors from each stage of the analysis. We demonstrated in an example that an integrated multistage framework can improve accuracy and precision of the estimates, compared to a hard cut-off two-step GSI approach. This method can be extended with an unlimited number of regional baselines if there are enough markers in a broad-scale baseline to allocate fish to broad-scale reporting groups. The regional baselines do not require any overlap in genetic markers with the broad-scale baseline or other regional baselines nor any overlap in the populations representing the broad-scale or regional baselines. These attributes made this approach much more cost-effective than past approaches where broad-scale baselines had to be updated with all critical regionally important markers.

The two-step analytic procedure was not a new concept. In the past, the two steps were conducted sequentially, and each as a separate process. At the first step of a GSI analysis, each individual in the mixture sample was assigned an identity with a level of certainty. Only individuals with assignment probabilities greater than a set threshold (e.g., 0.8) would proceed to the second step. In our analyses, we found that misassignments in the first step made the results at the second step unreliable. In our example, a 0.8 threshold on the assignment probability erroneously included some fish from Coastal Western Alaska and excluded more than half of Koyukuk and a good portion of Upper US and Tanana fish from the second step of the analysis (Figure 4). It introduced bias in the estimates. Lowering the threshold would include more middle/upper Yukon River fish, but it would risk falsely including more fish from Coastal Western Alaska. Most of it all, propagated errors were not accounted for in a hard cut-off two-step approach and credible intervals were underestimated (Figure 3).

Although we do not recommend a two-step approach, each stage of the integrated multistage framework can be done separately and combined at the end using conditional probability *p*(*Z*^(*regional*)^|*Z*^(*broad*)^) (Appendix B). This approach can be done using currently available GSI software without additional programming. However, this approach can only produce point estimates for reporting group proportions. Those who are mathematically inclined can obtain estimates for standard deviation by manually deriving the posterior distributions or using a parametric bootstrapping method. Still, with the development of Ms.GSI, one can measure uncertainties directly using the posterior output. Ms.GSI is arguably the most accurate and straightforward method among the available options. We also programmed Ms.GSI to provide assignment output for assessing individual identity.

The integrated multistage model was not impervious to biases either. Although we had shown that an integrated multistage model had lower RMSE and deviations compared to a hard cut-off two-step GSI approach, not all biases could be eliminated with an integrated multistage framework. Moran & Anderson (2018) developed a bootstrap procedure to reduce bias in the GSI estimates. A similar procedure may be developed for the integrated multistage framework to improve accuracy of the estimates.

## Conclusions

An integrated multistage framework provides superior estimates and measurement of uncertainty compared to the conventional hard cut-off two-step approach. It has the potential allowing researchers to conduct GSI for a wide geographic area without first developing a large-scale high-resolution genetic baseline or dividing mixture sample into smaller regions beforehand. Instead, researchers can utilize existing broad-scale baselines in combination with regional baselines. An integrated multistage framework that combines a broad-scale baseline with regional baselines may make collaborations easier between laboratories, institutions, or agencies from different regions. Most of all, it also means that a high-resolution baseline would only need to be developed regionally to focus on populations that were difficult to distinguish genetically. The results showed potentials for reduction of cost and time in development and improvement of genetic baselines.

## Acknowledgements

We thank the GCL staff for providing the Chinook baselines used in this analysis and for reviewing the earlier versions of this manuscript, Jim Jasper for his *R* code that provides the foundation for development of Ms.GSI, and Eric Anderson for his insights and comments on the conditional probability that forms the basis of the integrated multistage model.

## References

Ackerman, M. W., C. Habicht, and L. W. Seeb. 2011. Single-nucleotide polymorphisms (SNPs) under diversifying selection provide increased accuracy and precision in mixed-stock analyses of sockeye salmon from the Copper River, Alaska. Transactions of the American Fisheries Society. 140(3):865–881.

Barclay, A. W., D. F. Evenson, and C. Habicht. 2019. New genetic baseline for Upper Cook Inlet Chinook salmon allows for the identification of more stocks in mixed stock fisheries: 413 loci and 67 populations. Alaska Department of Fish and Game, Fishery Manuscript Series No. 19-06, Anchorage.

Beacham, T. D., J. R. Candy, B. McIntosh, C. MacConnachie, A. Tabata, K. Kaukinen, L. Deng, K. M. Miller, R. E. Withler, and N. Varnavskaya. 2005. Estimation of stock composition and individual identification of sockeye salmon on a Pacific Rim basis using microsatellite and major histocompatibility complex variation. Transactions of the American Fisheries Society. 134(5):1124–1146.

Beacham, T. D., J. R. Candy, C. Wallace, S. Urawa, S. Sato, N. V. Varnavskaya, K. D. Le, and M. Wetklo, 2009. Microsatellite stock identification of chum salmon on a Pacific Rim basis. North American Journal of Fisheries Management. 29(6):1757–1776.

Beacham, T. D., C. Wallace, K. Jonsen, B. McIntosh, J. R. Candy, E. B. Rondeau, J. S. Moore, L. Bernatchez, and R. E. Withler. 2020. Accurate estimation of conservation unit contribution to coho salmon mixed-stock fisheries in British Columbia, Canada, using direct DNA sequencing for single nucleotide polymorphisms. Canadian Journal of Fisheries and Aquatic Sciences. 77(8):1302–1315.

Beacham, T. D., C. Wallace, K. Jonsen, B. J. Sutherland, C. Gummer, and E. B. Rondeau. 2021. Estimation of conservation unit and population contribution to Chinook salmon mixed-stock fisheries in British Columbia, Canada, using direct DNA sequencing for single nucleotide polymorphisms. Canadian Journal of Fisheries and Aquatic Sciences. 78(10):1422–1434.

Brooks, S. P., and A. Gelman. 1998. General methods for monitoring convergence of iterative simulations. Journal of Computational and Graphical Statistics. 7:434–455.

Casella, G. and E.I. George. 1992. Explaining the Gibbs Sampler. The American Statistician. 46(3):167.

Ensing, D., W. W. Crozier, P. Boylan, N. O’Maoiléidigh, and P. McGinnity. 2013. An analysis of genetic stock identification on a small geographical scale using microsatellite markers, and its application in the management of a mixed-stock fishery for Atlantic salmon *Salmo salar* in Ireland. Journal of fish biology. 82(6):2080–2094.

Euclide, P. T., T. MacDougall, J. M. Robinson, M. D. Faust, C. C. Wilson, K. Y. Chen, E. A. Marschall, W. Larson, and S. Ludsin. 2021. Mixed-stock analysis using Rapture genotyping to evaluate stock-specific exploitation of a walleye population despite weak genetic structure. Evolutionary applications. 14(5):1403–1420.

Flannery, B. G., T. D. Beacham, J. R. Candy, R. R. Holder, G. F. Maschmann, E. J. Kretschmer, and J. K. Wenburg. 2010. Mixed-stock analysis of Yukon River chum salmon: Application and validation in a complex fishery. North American Journal of Fisheries Management. 30(5):1324–1338.

Gelman, A., and D. B. Rubin. 1992. Inference from iterative simulation using multiple sequences. Statistical Science. 7:457–472.

Gelman, A., J. Carlin, H. Stern, D. Dunson, A. Vehtari and D. Rubin. 2014. Bayesian Data Analysis, 3rd Edition. CRC Press. New York.

Gilbey, J., J. Coughlan, V. Wennevik, P. Prodöhl, J. R. Stevens, C. Garcia de Leaniz, D. Ensing, E. Cauwelier, C. Cherbonnel, S. Consuegra, and M. W. Coulson. 2018. A microsatellite baseline for genetic stock identification of European Atlantic salmon *(Salmo salar L.)*. ICES Journal of Marine Science, 75(2):662–674.

Howard, K. G., S. Garcia, J. Murphy, and T. H. Dann. 2019. Juvenile Chinook salmon abundance index and survey feasibility assessment in the Northern Bering Sea, 2014–2016. Alaska Department of Fish and Game, Fishery Data Series No. 19-04, Anchorage.

Lee, E., T. Dann, and H. Hoyt. 2021. Yukon River Chinook Genetic Baseline Improvements. Yukon River Panel Restoration and Enhancement Fund Final Report, URE-163-19N.

McKinney, G. J., C. E. Pascal, W. D. Templin, S. E. Gilk-Baumer, T. H. Dann, L. W. Seeb, and J. E. Seeb. 2020. Dense SNP panels resolve closely related Chinook salmon populations. Canadian Journal of Fisheries and Aquatic Sciences. 77(3):451–461.

Miller, J. A., M. R. Bellinger, J. T. Golden, L. Fujishin, and M. A. Banks. 2010. Integration of natural and artificial markers in a mixed stock analysis of Chinook salmon *(Oncorhynchus tshawytscha)*. Fisheries research. 102(1-2):152–159.

Moran, B. M. and E. C. Anderson. 2018. Bayesian inference from the conditional genetic stock identification model. Canadian Journal of Fisheries and Aquatic Sciences. 76(4):551–560. https://doi.org/10.1139/cjfas-2018-0016

Pella, J. and M. Masuda. 2001. Bayesian methods for analysis of stock mixtures from genetic characters. Fish. Bull. 99:151–167.

R Core Team (2022). R: A language and environment for statistical computing. R Foundation for Statistical Computing, Vienna, Austria. URL https://www.R-project.org/.

Samarasin, P., S. J. Latham, C. M. Guthrie III, and H. A. Araujo. 2019. What can We Learn About the Return Migration of Fraser River Sockeye Salmon from Catches in Alaska? North Pacific Anadromous Fish Commission, Technical Report No. 15: 40–43, 2019, doi: 10.23849/npafctr15/40.43.

Seeb, L. W., A. Antonovich, M. Banks, T. Beacham, R. Bellinger, S. Blankenship, M. Campbell, N. DeCovich, J. C. Garza, C. Guthrie, T. Lundrigan, P. Moran, S. Narum, J. Stephenson, J. Supernault, D. Teel, W. D. Templin, J. K. Wenburg, S. Young, and C. T. Smith. 2007. Development of a standardized DNA database for Chinook salmon. Fisheries. 32:540–552.

Stephenson J. J., M. R. Campbell, J. E. Hess, C. Kozfkay, A. P. Matala, M. V. McPhee, P. Moran, S. R. Narum, M. M. Paquin, O. S. Maureen, P. Small, D. M. Van Doornik, J. K. Wenburg. 2009. A centralized model for creating shared, standardized, microsatellite data that simplifies inter-laboratory calibration. Conserv Genet. 10:1145–1149. doi: 10.1007/s10592-008-9729-4.

Wirgin, I., L. Maceda, M. Tozer, J. Stabile, and J. Waldman. 2020. Atlantic coastwide population structure of striped bass *Morone saxatilis* using microsatellite DNA analysis. Fisheries research. 226:105506.

