## Appendix A for "Harnessing the power of regional baselines for broad-scale genetic stock identification: A multistage, integrated, and cost-effective approach"

Appendix A Table 1: Multistage model estimates for the Northern Bering Sea Chinook leave-one-out analysis with convergence diagnostics. $\hat{R}$ represents Gelman-Rubin potential scale reduction factor. $N_{eff}$ represents effective size of posterior samples.

| Group | Mean | Median | SD | Lower 90% CrI | Upper 90% CrI | $\hat{R}$ | $N_{eff}$ |
| --- | --- | --- | --- | --- | --- | --- | --- |
| Coastal W. Alaska | 0.29 | 0.29 | 0.05 | 0.21 | 0.37 | 1.00 | 1025.04 |
| Koyukuk | 0.08 | 0.08 | 0.02 | 0.05 | 0.13 | 1.00 | 4057.26 |
| Tanana | 0.20 | 0.20 | 0.03 | 0.14 | 0.26 | 1.00 | 2552.64 |
| Upper US | 0.13 | 0.13 | 0.03 | 0.09 | 0.18 | 1.00 | 6879.51 |
| Canada | 0.30 | 0.30 | 0.03 | 0.25 | 0.35 | 1.00 | 10051.76 |

Appendix A Table 2: Summary of model evaluation based on 50 leave-one-out analyses. ‘% Deviation > 0.1’ summarized 50 mean estimates of each model that deviate more than $\pm$ 0.1 from the true values for each reporting group. Root mean squared errors (RMSE) measure the sqare root of average squared errors of the reporting group proportions.

| Group | RMSE (Ms.GSI) | RMSE  (HC 2-step) | Bias (Ms.GSI) | Bias  (HC 2-step) | % Deviation (Ms.GSI) | % Deviation (HC 2-step) |
| --- | --- | --- | --- | --- | --- | --- |
| Coastal W. Alaska | 0.0526 | NA | 0.0257 | NA | 4 | NA |
| Koyukuk | 0.0415 | 0.0944 | -0.0244 | 0.0767 | 4 | 26 |
| Tanana | 0.0263 | 0.0357 | 0.0091 | 0.0289 | 0 | 0 |
| Upper US | 0.0164 | 0.0435 | -0.0038 | 0.0316 | 0 | 4 |
| Canada | 0.012 | 0.0666 | -0.0067 | 0.0541 | 0 | 10 |
