## Appendix B for "Harnessing the power of regional baselines for broad-scale genetic stock identification: A multistage, integrated, and cost-effective approach"

Another way to conduct GSI in an integrated multistage framework is by calculating conditional probability $P\left( Z^{\left( regional \right)}|Z^{\left( broad \right)} \right)$:

$$\begin{matrix} P & \left( Z_{k}^{\left( regional \right)}|Z^{\left( broad \right)} \right)= \\ & \sum_{m=1}^{M} \frac{P\left( Z_{m}^{\left( broad \right)}|Z_{m,k}^{\left( regional \right)} \right)\cdot P\left( Z_{m,k}^{\left( regional \right)} \right)}{\sum_{k=1}^{K} P\left( Z_{m}^{\left( broad \right)}|Z_{m,k}^{\left( regional \right)} \right)\cdot P\left( Z_{m,k}^{\left( regional \right)} \right)}=\frac{\sum_{m=1}^{M} P\left( Z_{m}^{\left( broad \right)} \right)\cdot P\left( Z_{m,k}^{\left( regional \right)} \right)}{\sum_{m=1}^{M} P\left( Z_{m}^{\left( broad \right)} \right)} \end{matrix}$$

(Eq. 2)

In this approach, we would conduct GSI for each baseline separately. First, use the broad-scale baseline to assign each fish in the mixture data to the broad-scale groups. For each fish $m$, $\rho_{m}|z_{m}^{\left( broad \right)}$ is the posterior probability of being from a regional group based on the broad-scale markers. $\rho_{m}|z_{m}^{\left( broad \right)}$ can be derived by summarizing the proportion of group assignments that are within the regional baseline in the posterior output for fish $m$. Next, conduct GSI using the regional baseline. For each fish $m$, posterior probabilities for assignment to regional groups are vectors $\pi_{m}|z_{m}^{\left( regional \right)}$ and can be derived by summarizing the group proportions in the posterior output for individual assignments. Lastly, calculate a point estimate of the proportion of fish from each regional population by summing $\rho_{m}\left| z_{m}^{\left( broad \right)}\cdot\pi_{m} \right|z_{m}^{\left( regional \right)}$ over all fish in the mixture sample and then normalizing with $\sum_{m=1}^{M} \rho_{m}|z_{m}^{\left( broad \right)}$. There would be no direct measurements of uncertainty using this approach.
